# Light-induced polyp retraction and tissue rupture in the photosensitive, reef-building coral *Acropora muricata*

**DOI:** 10.1101/862045

**Authors:** Pierre Philippe Laissue, Yan Gu, Chen Qian, David J. Smith

## Abstract

Coral reefs are in alarming decline due to climate emergency, pollution and other man-made disturbances. The numerous ecosystem services derived from coral reefs are underpinned by the growth and physical complexity of reef-forming corals. Our knowledge of their fundamental biology is limited by available technology. We need a better understanding of larval settlement and development, skeletogenesis, interactions with pathogens and symbionts, and how this biology interacts with environmental factors such as light exposure, temperature, and ocean acidification. We here focus on a fast-growing key coloniser, *Acropora muricata*. To enable dynamic imaging of the photosensitive organism at different scales, we developed light-sheet illumination for fluorescence microscopy of small coral colonies. Our approach reveals live polyps in previously unseen detail. An imaging range for *Acropora muricata* with no measurable photodamage is defined based upon polyp expansion, coral tissue reaction, and photobleaching. We quantify polyp retraction as a photosensitive behavioural response and show sparse zooxanthellar expulsion and coral tissue rupture at higher intensities of blue light. The simple and flexible technique enables non-invasive continuous dynamic imaging of highly photosensitive organisms with sizes between 1 mm^3^ and 5 cm^3^, for eight hours, at high temporal resolution, on a scale from multiple polyps down to cellular resolution. This live imaging tool opens a new window into the dynamics of reef-building corals.

## Introduction

Live imaging is a potent approach for the investigation of fundamental processes and structures of reef-building corals at microscopic level. Many microscopy-based studies of corals rely on techniques imaging the calcareous skeleton from which the tissue was removed, or using fixed samples of decalcified coral tissue. Therefore, the complex three-dimensional interactions of coral tissue, endosymbiontic algae and aragonite skeleton have been little studied at the tissue and cellular level. Knowledge of such interactions can greatly add to our understanding of fundamental coral biology which is required to improve strategies for the conservation of coral reefs. In particular, fluorescence microscopy can reveal a lot, since it opens up the possibility to use coral tissue autofluorescence as a non-invasive, intrinsic marker of health and disease (Caldwell et al., 2017; Kenkel et al., 2011) and the monitoring of chlorophyll autofluorescence of the photosynthetic symbionts embedded in the coral tissue (Mullen et al., 2016; Shapiro et al., 2016). Fluorescent dyes can also be used in live imaging (Neder et al., 2019; Ohno et al., 2017; Venn et al., 2009).

So far, only conventional techniques have been used to image live coral fluorescence on the cellular scale. Confocal laser scanning microscopy (CLSM) of live polyps has been used in *Montipora capitata* at different spatial scales, to characterize overall diversity of natural fluorescence and to spatially localize the arrangement of fluorescent pigments (Caldwell et al., 2017). Inverted microscopy from below has been used to image several microns deep into the calcifying layers (Neder et al., 2019; Ohno et al., 2017; Shapiro et al., 2016; Venn et al., 2011). The approach works well when the main interest lies in the calcification of the flat bottom layer. However, this approach is not well suited for large samples with complex three-dimensional growth such as small coral colonies. Compared to model organisms commonly used in fluorescence microscopy, coral colonies with multiple polyps are considerably larger (Fig. 1 A). We chose light-sheet fluorescence microscopy, also called Selective Plane Illumination Microscopy (SPIM, Huisken et al., 2004), a versatile technique for samples that cannot be mounted between glass (Gutiérrez-Heredia et al., 2012). For the large coral samples, we created a very wide light-sheet to capture a large field of view (FOV). The common OpenSPIM-type static Gaussian light-sheet (Pitrone et al., 2013) is around one millimetre wide. It thus reveals only a small part of a coral colony (Fig. 1 B, left side, short blue bar). By comparison, as we sweep the beam laterally, our light-sheet can be made as wide as two centimetres (Fig. 1 B, right side, wide blue bar). We thus call it the large selective plane illuminator (L-SPI).

The key strength of light-sheet fluorescence microscopy is the ability to minimise photodamage during live imaging. This was an essential consideration for observing the photosensitive species *Acropora muricata*. In conventional microscopy, fluorescence in tissue, cells or aragonite skeleton is excited and detected through the same (inverted) objective (Fig. 1 C, ‘conventional’). The consequence is that for each focal plane, the entire sample is illuminated. This makes conventional fluorescence microscopy techniques not well suited for long-term imaging of photosensitive samples. By contrast, the excitation pathway in light-sheet microscopy is uncoupled and comes in from the side through a cylindrical lens (Fig. 1 C, ‘L-SPI’). It illuminates only the focal plane. This enables live imaging in three spatial dimensions at minimal light exposure.

**Fig. 1.**
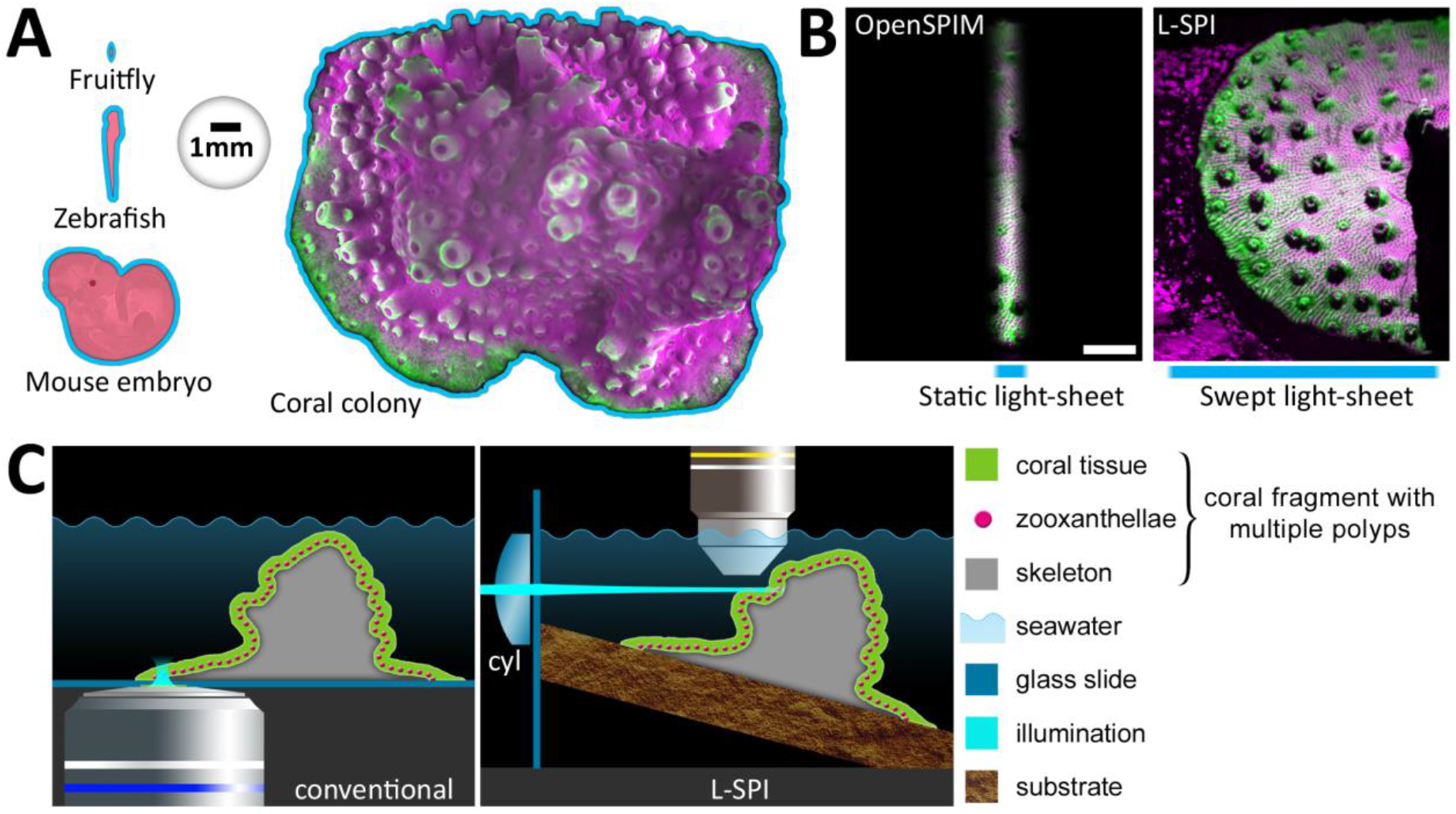
A) Size of commonly used samples and of a small colony of reef-building corals. B) Left side: Width of an OpenSPIM-type static light-sheet. The scale bar is 2 mm. Right side: Our approach (L-SPI) generates a much wider light-sheet, which allows illuminating large parts of a small coral colony in a single FOV. C) Left side: The conventional approach uses inverted microscopy to excite and detect fluorescence in a colony’s growing edge from below. Right side: Our approach (L-SPI) uses light-sheet excitation created by a swept laser beam entering from the left side and passing through a cylindrical lens (‘cyl’), with fluorescence emission detected from above. This enables low-light imaging of complex topology or individual polyps.

We used small colonies of *Acropora muricata* (Linné, 1758) as an experimental model for coral development and behaviour. Corals of the genus *Acropora* are fast-growing, ecologically crucial key architects of coral reef ecosystems, greatly contributing to their complex three-dimensional structure (Renema et al., 2016). They can also recover rapidly from environmental disturbances, making them important re-colonisers (Sweatman et al., 2011). Rather than using cell or tissue culture-based samples (Mass et al., 2012), single primary polyps (Neder et al., 2019; Ohno et al., 2017) or single polyps obtained through bail-out techniques (Shapiro et al., 2016), we here use small coral colonies with multiple polyps and a shared gastrovascular cavity. They allow the study of systemic responses in an unstressed, fully established colony. Our non-invasive observation technique expands the live-imaging toolbox to photosensitive species and long-term observation of chlorophyll fluorescence in algal symbionts.

## RESULTS

### Morphology of large *Acropora formosa* colonies at multiple scales

Width and thickness (waist) of the L-SPI light-sheets were adapted to cover samples of different sizes. This enabled imaging of a coral colony at multiple scales (Fig. 2). Wide light-sheets (2 cm width) with a large waist (20.7 ± 0.8 μm) were used to provide an overview of large parts of a colony in a single field-of-view (FOV). Figure 2 A shows multiple polyps in a FOV of 26 mm x 19 mm. These datasets are acquired in three spatial dimensions (x, y and z), also called z-stacks. They reveal the topology of the cup-shaped corallites. A corallite is the protective, skeletal cover into which a single polyp can retract (Gladfeiter, 1982; Gladfelter, 1983; Gladfelter, 2007). Corallites are very flat at the growing edge, and rise up with increased distance from the edge (which correlates with their advanced developmental stage). Since the coral skeleton is entirely opaque, a certain amount of shadowing cannot be avoided. Note that all polyps have emerged. The different lateral resolutions and light-sheet dimensions are summarised in Supplementary Table 1.

Using a light-sheet with thinner waist (11.7 ± 0.8 μm) allows focussing on a single coral polyp and large parts of its surrounding tissue (Fig. 2 B). The coral tissue has strong green autofluorescence and a fibrous structure. It clings to the side of the prominent skeletal spines (Gladfeiter, 1982; Gladfelter, 1983; Gladfelter, 2007). Zooxanthellae are arranged in wavy bands within the tissue. At this magnification, single zooxanthellae in the polyp tentacles are resolved (Fig. 2 B i). This enables automated counting using simple image processing (Fig. 2 B ii). A small polyp in an earlier stage of development is shown at higher resolution in Fig. 2 C (supplementary movie S1). Subcellular structures of zooxanthellae are visible at this scale. Zooxanthellae have a diameter of 8.5 ± 1.0 μm (n = 203). They are densely packed in the budding tentacles of the polyp, arranged as a single-cell layer on each side of a tentacle. In mature polyps, these layers often join up to form a funnel-shaped single-cell layer of zooxanthellae.

**Fig. 2.**
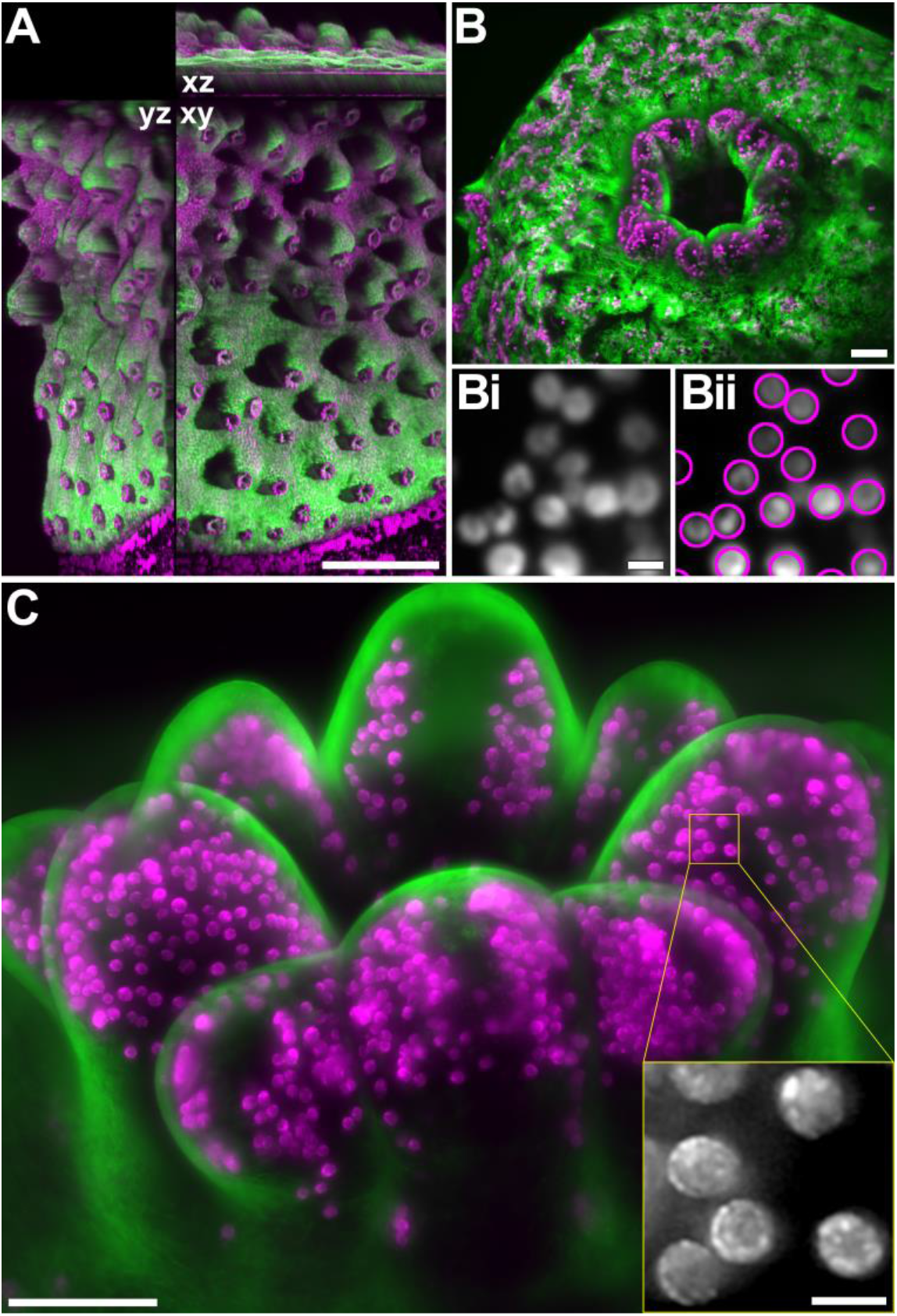
Volumes of photosensitive *Acropora muricata* obtained with the L-SPI. In all images, coral tissue autofluorescence is green, while magenta shows the chlorophyll autofluorescence of the symbiotic zooxanthellae. A) Large field of view (FOV) of multiple polyps in a coral colony. The orthogonal sections (xy, xz and yz) show the topology of its growth. Scalebar 5 mm. B) Larger magnification (10x) of a coral polyp. Scalebar 100 μm. Bi) Close-up of a small region showing individual zooxanthellae. Scalebar 10 μm. Bii) Automated identification of individual zooxanthellae. C) A small, developing coral polyp (imaged at 20x) showing the distribution of symbiotic algae embedded in the tentacles. Scalebar 100 μm. Inset: High magnification view of zooxanthellae showing subcellular detail. Scalebar 10 μm.

### The rate of polyp expansion depends on excitation light irradiance in *Acropora formosa*

We have previously suggested that the expansion and contraction of polyps can be used to determine physiological and excessive levels of excitation light used for microscopic observation of *Acropora muricata* (Laissue et al., 2017), and quantify it in this study. Polyps were retracted at the start of an imaging experiment after transferring the coral fragment from the main aquarium to the observation vessel. As shown in Fig. 3, connecting the tips of a polyp’s tentacles was used to determine the area of expansion. A normalised value of one means full expansion (or highest expansion for the duration of observation), and zero denotes full contraction (or lowest expansion for the duration of observation). At an average illumination power of 20 μW (Fig. 3 A, top row; supplementary movie S2), corresponding to an irradiance of 18.4 mW/cm^2^, polyps had expanded by 84% within 90 minutes (median value; 95% median confidence interval 0.75 – 0.90). Within the same timeframe, but at 90 μW excitation power (82.6 mW/cm^2^ irradiance), expansion was less than half (41%; 95% confidence interval 0.14 – 0.64; p = 0.012 using randomisation test (Hooton, 1991; Nuzzo, 2017) (Fig. 3 A, bottom row; supplementary movie S3)). Conversely, polyps which had expanded at low excitation irradiance (9.2 mW/cm^2^) would retract at high irradiance (59.7 mW/cm^2^) (Fig. 3 B; supplementary movie S4). This provided an initial estimate of tolerable versus excessive levels of blue light irradiance for fluorescence excitation. The different illumination conditions used in this study are summarised in Supplementary Table 2.

**Fig. 3.**
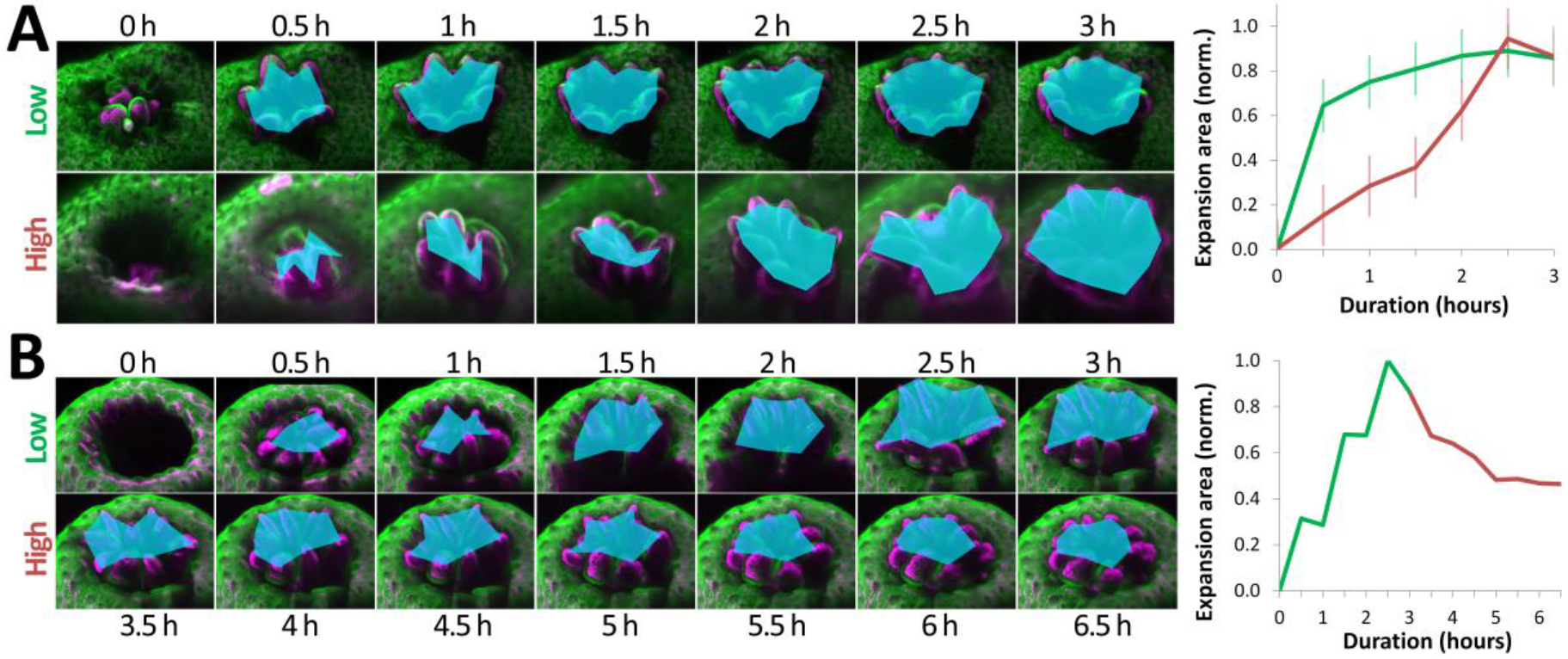
Polyp emergence and retraction under low and high irradiance. A) Images of two polyps in low and high illumination conditions. Emergence is quantified using the area determined by connecting the tips of tentacles. At low irradiance (18.4 mW/cm^2^), polyps have nearly fully emerged after one hour. At high irradiance (82.6 mW/cm^2^), polyp emergence takes more than twice as long. This is quantified in the line graph on the right. Error bars show standard error. B) Images of a polyp switching from low to high irradiation. The polyp emerges in low light (9.2 mW/cm^2^) and retracts in high light (59.7 mW/cm^2^). This is quantified in the line graph on the right.

### Non-invasive continuous imaging for at least six hours does not cause light-induced stress

Using the light irradiance ranges which allow rapid polyp expansion, we investigated longer exposure times to determine other indicators of stress and potential photodamage. Long time-lapse recordings of sequential z-stacks were taken therefore. Figure 4 (supplementary movie S5) shows the typical dynamics of a coral polyp of *Acropora muricata* over six hours of continuous imaging at low light. The polyp expanded rapidly within half an hour, and stayed expanded for the rest of the time-lapse acquisition. There was no change in the number and distribution of zooxanthellae (Fig. 4 C, supplementary movie S6). Since fluorescence intensity and photobleaching correlate with the generation of reactive oxygen species, photobleaching is a useful proxy for the semiquantitative assessment of phototoxicity (Carlton et al., 2010; Laissue et al., 2017; Reynaud et al., 2008). Here, photobleaching of the coral tissue was minimal (10% reduction in fluorescence intensity) and reached a plateau after one hour (see also Fig. 7 C). We hypothesised that these various characteristics signified the absence of light-induced stress. All colonies imaged this way were re-used for imaging and showed no signs of immediate or delayed photodamage. A median value of eight hours continuous image acquisition was achieved (n=6) using a power of 10 μW (9.2 mW/cm^2^ irradiance). The average exposure was 184 ± 50 mJ. Given a typical polyp size of 1 mm x 1 mm x 0.8 mm, this results in a very low energy density of 0.23 nJ/μm^3^. With an average of 220 volumes over an 8 h period of observation, this implies an exposure of 836 μJ per stack and an energy density of 1.0 pJ/μm^3^ per stack.

**Fig. 4.**
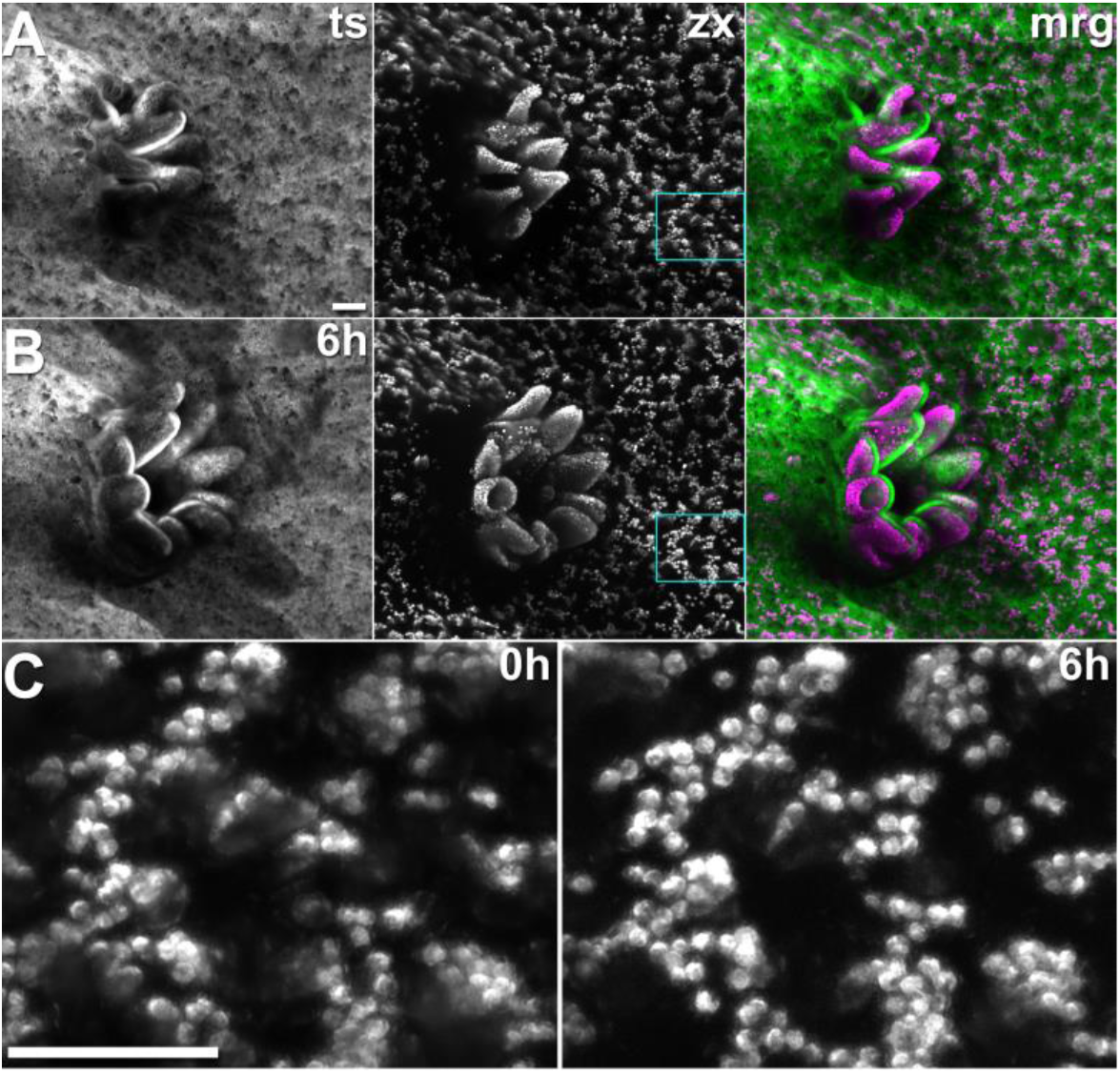
Still frames from sequential z-stacks recorded over six hours at low light (9.2 mW/cm^2^). A) Polyp at start of acquisition. From left to right: Coral tissue autofluorescence (ts), chlorophyll autofluorescence of the zooxanthellae (zx), and merged image (tissue green, zooxanthellae magenta). B) The same polyp after six hours of continuous image acquisition. C) Close-up of the cyan boxes in A and B showing the unchanged arrangement of the zooxanthellae inside the tissue. Scalebars 100 μm.

### Increased irradiance provokes sparse zooxanthellar expulsion

We first imaged a dynamic polyp over seven hours of continuous acquisition in low light (9.2 mW/cm^2^, Fig. 5 A, B). It expanded fully, to the point where it left the FOV. Sample and perfusion were manually adjusted after five hours, causing the polyp to contract and expand again. At seven hours, close-ups of the zooxanthellar bands showed densely packed zooxanthellae (Fig. 5 C, left side; supplementary movie S7). For the following 90 minutes, the laser excitation power was increased from 10 μW to 65 μW (9.2 mW/cm^2^ and 59.7 mW/cm^2^, respectively). This caused the polyp to contract. Another sign of light-induced stress was the expulsion of single zooxanthellae. Dark holes are visible where single zooxanthellae have been expelled (Fig. 5 C, see white arrowheads on right side; supplementary movie S7).

**Fig. 5.**
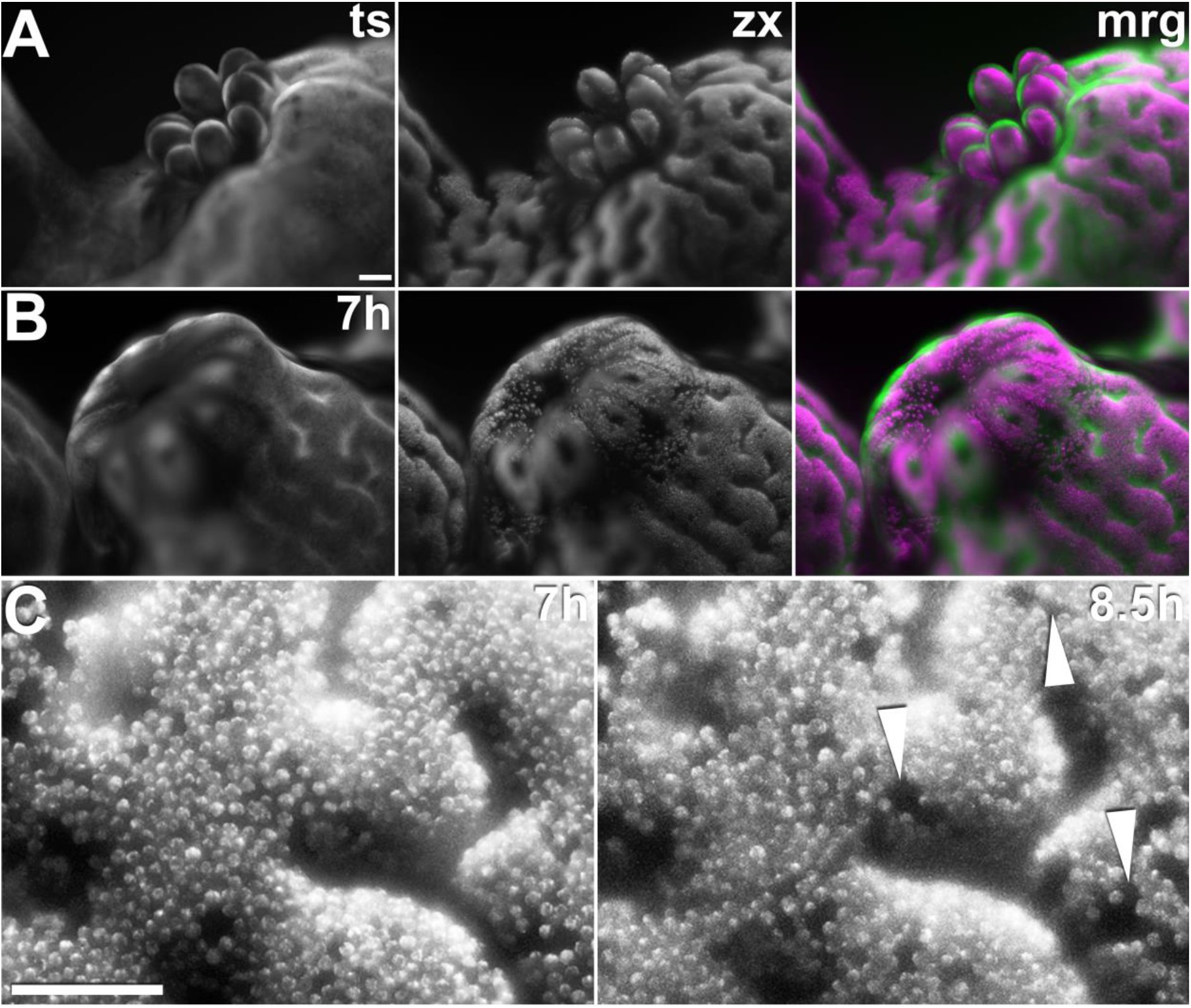
Still frames from sequential z-stacks recorded over eight and a half hours. A) Polyp at start of acquisition. From left to right: Coral tissue autofluorescence (ts), chlorophyll autofluorescence of the zooxanthellae (zx), and merged image (mrg; tissue green, zooxanthellae magenta). Scale bar 100 μm. B) The same polyp after seven hours of continuous image acquisition at low light (9.2 mW/cm^2^). Scale bar 100 μm. C) Close-up of B (zooxanthellae) after seven hours of low-light illumination (left), followed by 90 minutes at high light (right, 59.7 mW/cm^2^). Holes (white arrowheads) mark sites of expelled zooxanthellae. Scale bar 100 μm.

### High irradiance leads to tissue rupture

Photodamage was far more severe when a high excitation power was used from the start. Figure 6 (supplementary movie S8) shows that this increased excitation power eventually caused tissue rupture. Using 52 μW illumination (47.8 mW/cm^2^ irradiance) for continuous image acquisition, no adverse reactions were observed for the first five hours, apart from pronounced photobleaching of the coral tissue. After seven hours, a surge in coral tissue and chlorophyll autofluorescence indicated a strong reaction to the high light. This was followed by tissue contraction and rupture, starting at nine hours. The illuminated area was irreversibly damaged and the entire colony perished within two days.

**Fig. 6.**
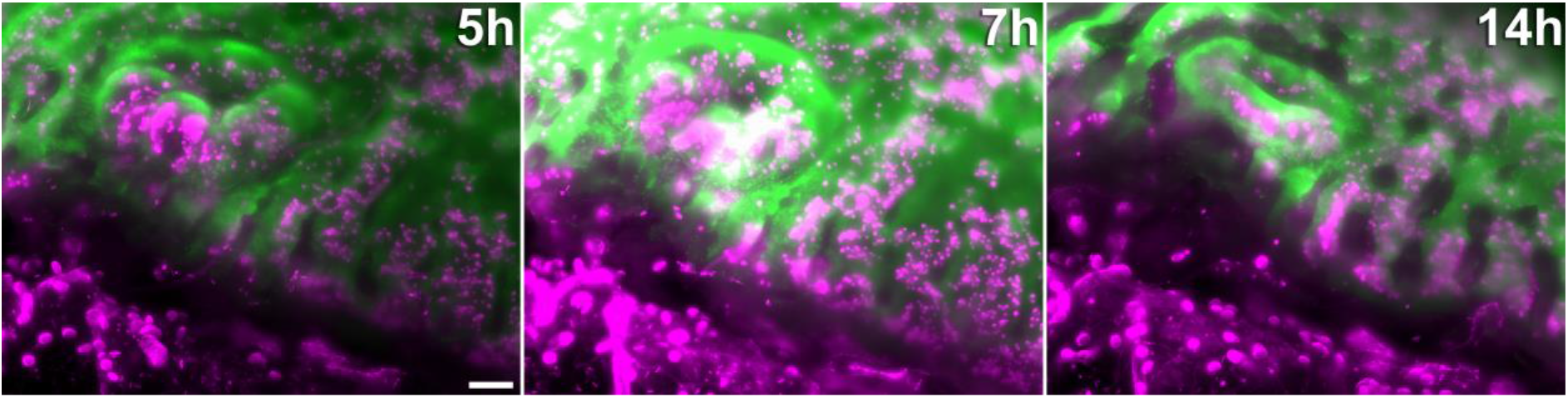
Still frames from a 16 hour continuous recording in excessive light conditions *(47.8 mW/cm^2^)*. Coral tissue autofluorescence (green) and chlorophyll autofluorescence (magenta) are shown after five hours of continuous imaging (left). At seven hours (middle), a surge in autofluorescence intensity was visible. This was followed by tissue rupture (occurring between nine and eleven hours), which lead to the growing edge dissolving after 14 h (right). Scale bar 100 μm.

### Conventional fluorescence microscopy methods cause more photobleaching and photodamage compared to the L-SPI

Using confocal laser scanning microscopy (CLSM) for image acquisition of *Acropora muricata* in three spatial dimensions invariably caused polyps to retract deeply into their corallites, even at a very low excitation power of 1.1 μW (Fig. 7 A, supplementary movie S9). Observing polyp dynamics in *Acropora muricata* was not possible using CLSM. Attempting longer-term time-lapse recordings at the growing edge resulted in marked tissue contraction and eventual rupture (Fig. 7 B). This is not surprising since CLSM produces a focal laser point with very high irradiance: Using a 10x lens with a numerical aperture (NA) of 0.3 and very low laser excitation (1.1 μW) results in 34’414 mW/cm^2^, which is larger by a factor of 3’741 compared to the light-sheet’s irradiance at the non-invasive level (9.2 mW/cm^2^). In conclusion, in the case of *Acropora muricata*, CLSM provided live morphology in three spatial dimensions (3D) due to its optical sectioning capability, but the inflicted photodamage limited acquisition to a single timepoint.

We then compared the L-SPI to widefield epifluorescence imaging (WFM). We wanted to see if WFM might be able to perform as well as light-sheet microscopy purely in terms of photobleaching. To reduce light exposure as much as possible for WFM, we used lower magnification (CFI Plan Apochromat λ 4X, NA 0.2; Nikon Corporation, Tokyo, Japan), enabling large 50 μm z-steps, so that a polyp was covered using only 11 to 13 z-planes. We further limited acquisition of these z-stacks to 5 minutes intervals, resulting in longer dark phases to further minimise photobleaching. At 1.0 mW/cm^2^, irradiance was ten times lower compared to LSFM. However, despite these efforts to minimise exposure in WFM, photobleaching of the coral tissue was higher compared to the L-SPI (Fig. 7 B, supplementary movie S1). The L-SPI covered six times more z-planes, nearly two and a half times higher time resolution, and at least one and a half times higher lateral resolution. Thus, the L-SPI, compared to sparse WFM, provides over 22 times more information while still causing 15% less photobleaching after six hours (86% and 71% of initial fluorescence intensity, respectively).

**Fig. 7.**
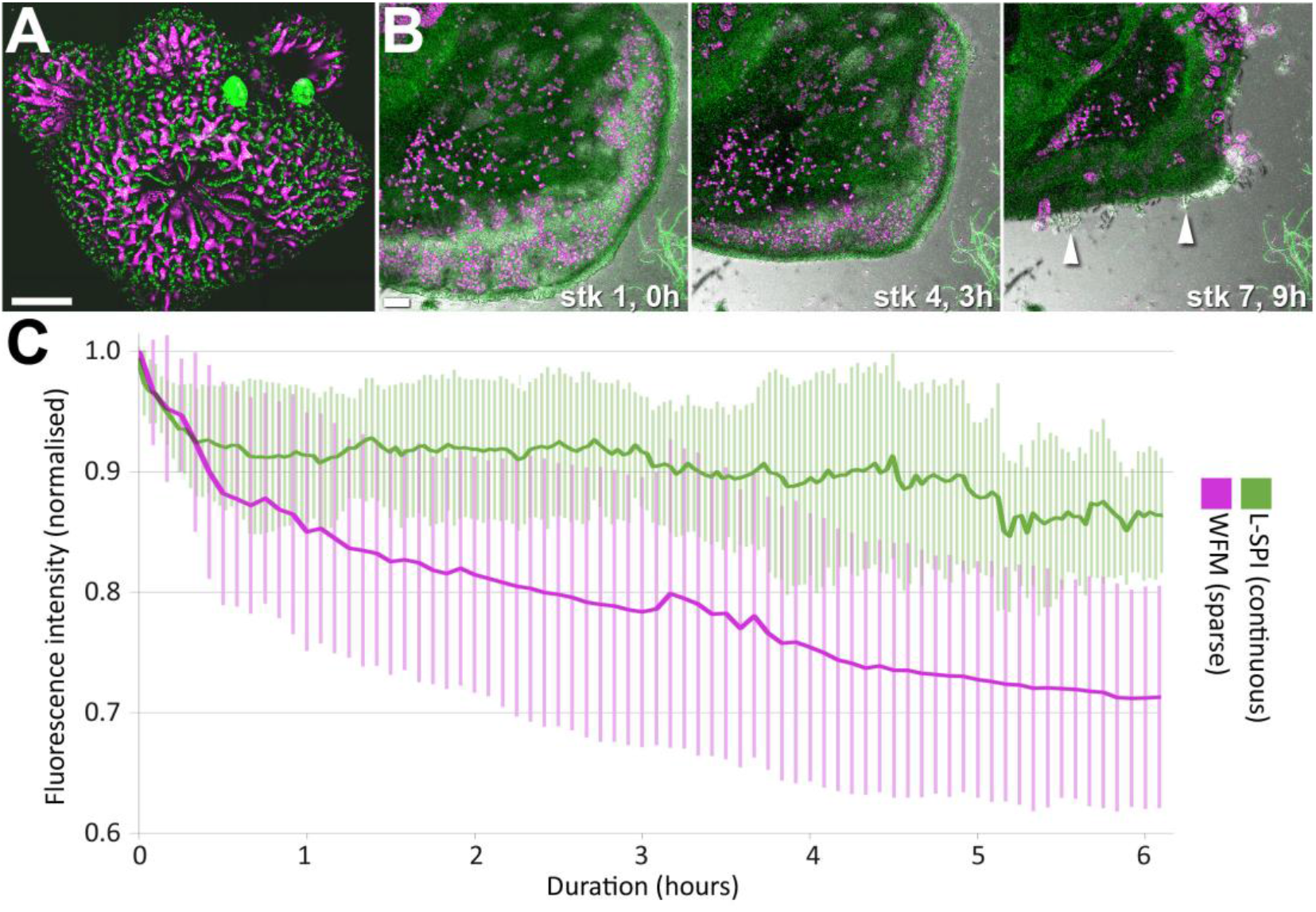
A) Polyps at the tip of a coral branch imaged with confocal laser scanning microscopy (CLSM). The large terminal polyp in the image centre, and the three smaller polyps in the upper half of the image, had all fully retracted into their corallites. Scalebar 500 μm. B) Severe retraction and rupture of coral tissue at the growing edge in *A. muricata*. At timepoint 0 in the first z-stack (stk 1, 0h), the tissue of the growing edge was expanded. Three z-stacks and three hours later (stk 4, 3h), the edge had retracted. Scans were then further reduced to one scan every two hours. However, after seven stacks (stk 7, 9h), the tissue started to rupture, exposing skeletal elements (white arrowheads) and compressing zooxanthellae in the gastrovascular cavity. Scalebar 100 μm. C) Comparison of photobleaching using sparse WFM (magenta) and L-SPI (green). Despite low irradiance (1.0 mW/cm^2^) and reduced volumetric and temporal imaging for WFM, photobleaching was 15% higher after six hours compared to the L-SPI.

## DISCUSSION

We have developed and applied a novel light-sheet approach to the study of reef-building corals. The instrument is capable of long-term three-dimensional imaging in photosensitive *Acropora muricata* without any measurable adverse reactions. This allowed us to observe the dynamics of small coral colonies in unprecedented temporal, spatial and spectral detail. We depict zooxanthellae arranged in bands within the coral tissue, and as a dense single layer in polyp tentacles. We demonstrate that polyp retraction is a photosensitive response, and define a non-invasive range for fluorescence excitation. A median value of 8 hours (mean value 8.2 ± 2.0 h) continuous image acquisition was possible using 488 nm excitation at a power of 10 μW (9.2 mW/cm^2^ irradiance). This resulted in an average exposure of 184 ± 50 mJ, with coral colonies showing no signs of short-term light-induced stress or long-term photodamage. Just outside of this non-invasive range, sparse expulsion of zooxanthellae was an early sign of stress. Substantially exceeding the non-invasive level lead to contraction and rupture of the coral tissue.

The L-SPI performed better than the conventional fluorescence microscopy methods WFM and CLSM. Despite using low excitation power, *Acropora muricata* was particularly susceptible to CLSM. Quantitative comparisons to previous studies using conventional microscopy (WFM, CLSM and spinning disk confocal) for live imaging of *Montipora capitata* (Caldwell et al., 2017; Nielsen et al., 2018), *Acropora digitifera* (Ohno et al., 2017), *Pocillopora damicornis* and *Stylophora pistillata* (Mullen et al., 2016; Neder et al., 2019; Shapiro et al., 2016; Venn et al., 2009; Venn et al., 2011; Venn et al., 2013) was not possible as no power measurements were reported. Substantial differences in phototolerance may exist between different species.

Polyp expansion and retraction is a fundamental behaviour in cnidarians (Swain et al., 2015). During daytime, branched corals with small polyps expand their tentacles to expose photosynthetic symbionts, localized within the tentacles, to light (Levy et al., 2003; Levy et al., 2006; Sebens and DeRiemer, 1977). Conversely, they retract their tentacles in high light conditions to protect the photosynthetic symbionts from damaging irradiance (Brown et al., 1994; Crossland and Barnes, 1977; Horridge, 1957; Salih et al., 2000). Maximum photobehaviour response has been demonstrated in the blue/green zone (Gorbunov and Falkowski, 2002; Levy et al., 2003). Our observations on photosensitive polyp dynamics are consistent with these published findings. Levy and coworkers (Levy, 2003) report average values at 5 m depth in Gulf of Eilat of around 12 mW/cm^2^ from about 11 am to 4 pm. Our non-invasive imaging range (9.2 mW/cm^2^ (375 μmol/s/m^2^) for an average of 8.3 ± 2.0 hours) broadly agrees with this value. It should also be noted that the 9.2 mW/cm^2^ are peak values, not average values, for the region illuminated (between 80 ms and 1 s per image) by the light-sheet. The light-sheet passing over the coral is not dissimilar to the effect of transient peak values of sunlight from water lensing, caused by ripples and waves at the top of the water.

Upon settling at the beginning of an imaging session, coral tissue often appears to swell within the first ten to twenty minutes. This is likely caused by influx of seawater into the gastrovascular cavity. This tissue inflation also appears to contribute to a drop in autofluorescence intensity. Thus, the drop in coral tissue autofluorescence observed at low light levels using the light-sheet may be only partly due to actual photobleaching.

The highly photosensitive *Acropora muricata* proved to be an ideal test-bed for live imaging at very low light exposure. Only autofluorescence was used for imaging, which also meant that, under stringent environmental control (temperature and perfused seawater), any signs of stress could be attributed to the illumination conditions alone.

The instrumental setup for the L-SPI is simple and flexible. It performs best with highly photosensitive organisms, and can image small to large sample volumes (1 mm^3^ to 5 cm^3^). These attributes enable studying the responses of tissue and symbionts on different spatial scales, at high temporal resolution and over several hours. This live imaging tool will continue to provide a new window into the dynamics of reef-building corals.

## MATERIALS & METHODS

### Sample preparation

Large coral colonies of *Acropora muricata* (Dana, 1846) were fragmented into smaller colonies (around 2 cm x 2 cm x 1 cm) with multiple polyps and grown on various substrates pre-treated with crustose coralline algae using veterinary-grade cyanoacrylate (superglue) (Reynaud-Vaganay et al., 1999; Shafir et al., 2001; Shafir et al., 2003) or fixed directly to the imaging chamber using inert putty. Imaging chambers were assembled from large optical-grade microscope slides (Agar Scientific, Elektron Technology Ltd, Essex, UK) using fungicide- and solvent-free silicon (Everbuild Building Products Ltd., Leeds, UK). The small aquaria were perfused at 27°C using a peristaltic pump.

### Large selective plane illumination

The L-SPI setup for coral *in vivo* imaging is detailed below (Fig. 8). Components and requisite weblinks are listed in Supplementary Table 3. The instrument was originally based on the OpenSPIM design (Pitrone et al., 2013). A 488 nm Optically Pumped Semiconductor Laser (OBIS 488 nm LS 100 mW, Coherent, Inc., Santa Clara, CA, USA) was controlled using an Arduino microcomputer and the open source software μManager (Edelstein et al., 2010; Stuurman and Swedlow, 2012). The beam was expanded twice and reflected onto a rotating mirror (AN8248NSB, Panasonic Corporation, Osaka, Japan) extracted from an old laser printer (for an equivalent rotating mirror, see Supplementary Table 3). The robust brushless motor was powered by a 375W linear DC variable voltage power supply (Maplin Electronics, Wombwell, UK). The rotating mirror fans the beam out laterally, creating a large light-sheet. Its width was controlled using an adjustable mechanical slit. A 50/50 beamsplitter combined with a prism split the light sheet in opposite directions. After each arm passed through a cylindrical lens, the two light sheets were recombined at a right angle, to reduce shadowing without the need to rotate the sample. Small coral colonies were individually stepped through the light-sheets using a motorized translation stage (Thorlabs Inc., Newton, New Jersey, USA).

Blueprint, components and weblinks required to build the instrument presented are provided in this study. The L-SPI has also been developed into a commercial version (Cairn Research Ltd., UK) to allow researchers who do not have the time or expertise to build the instrument described to benefit from its advantages.

**Fig. 8.**
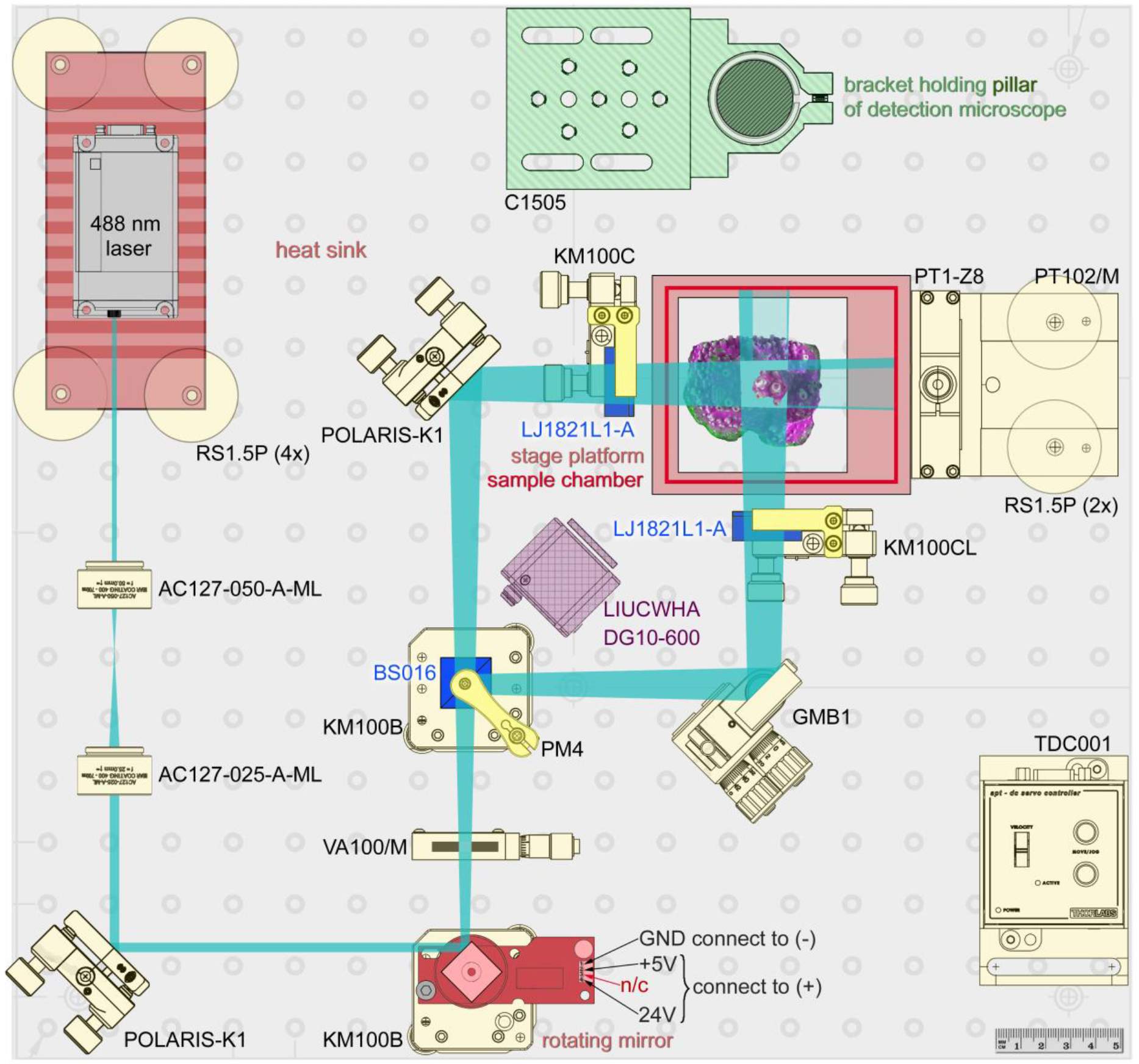
Schematic view of the illumination path for generating two broad light-sheets, as viewed from above. Briefly, the laser beam (top left) is expanded twice and projected onto the rotating mirror, which fans out the beam into a wide light-sheet. The light-sheet passes through a 50/50 non-polarising beamsplitter, generating two light-sheets of identical dimensions. The coral sample is then stepped through these light-sheets while images are acquired from above using any upright micro- or macroscope. The components list is provided in Supplementary Table 3.

### Detection

Images are taken from above using any upright micro- or macroscope. We have used two main approaches: A compound microscope with low (2x-5x) to medium (10x-40x) magnification objectives, and a macroscope (or stereomicroscope) with low magnification objectives (1x-2x) and zoom function. Images with large fields of view were taken using an SMZ25 stereomicroscope (Nikon Corporation, Tokyo, Japan), equipped with P2-SHR Plan Apochromatic 1x (NA 0.156, WD 60 mm) and 2x (NA 0.312, WD 20 mm) objectives. A Retiga 6000 CCD camera (Photometrics, AZ, USA) was used for detection, combined with an OptoSpin filter wheel (Cairn Research, Faversham, UK). Motorised z-stage and cameras were controlled using the open-source software μManager (Edelstein et al., 2010). Higher-resolution images of single polyps were taken using a BX41 compound microscope with water immersion objectives. A table of the different objectives used is provided in Supplementary Table 2.

### CLSM and WFM

A Nikon A1si CLSM (Nikon Corporation, Tokyo, Japan) with a spectral detector unit was used to define the autofluorescence signatures of coral tissue and zooxanthellae. Using an Argon-Ion laser at 488 nm (40 mW, Melles Griot plc, Carlsbad, CA, USA), coral tissue autofluorescence had its emission maximum at 500 nm and was collected using a 525/50 nm filter, while zooxanthellae showed characteristic chlorophyll *a* emission at a maximum of 685 nm, acquired by using a 700/75 nm emission filter. Objectives were CFI Plan Apochromat λ 4X (NA 0.2, WD 20 mm) and CFI PlanFluor 10x (NA 0.3, WD 16 mm). Images were acquired in two channels using one-way sequential line scans. NIS-Elements (version 3.21.03, build 705 LO) was used to acquire images.

WFM was performed using a Nikon Eclipse Ti-E main body and NIS-Elements (version 3.21.03, build 705 LO) for image acquisition. Objectives were a CFI Plan Apochromat λ 4X (NA 0.2, WD 20 mm) and a CFI PlanFluor 10x (NA 0.3, WD 16 mm). Excitation wavelength was 470 ± 10 nm using a pE-2 LED illuminator (CoolLED Ltd., Andover, UK). Hard-coated interference filters (Semrock Inc., IDEX corp., IL, USA) were used. For chlorophyll autofluorescence, these were a chromatic reflector at 665 nm and a long-pass emission filter at 664 nm. For tissue autofluorescence, we used a chromatic reflector at 505 nm and a single band emission filter at 535/40 (Chroma Technology Corp., VT, USA). Images were acquired with a Retiga 6000 CCD camera (Photometrics, AZ, USA). Power measurements for all systems were done with an ML9002A optical handy power meter (Anritsu Corp., Japan) measured at the sample.

### Image acquisition and processing software

Image processing was done using the open-source software FIJI (Schindelin et al., 2012). As the L-SPI takes images from a single viewpoint, no fusion of multiple views was required. We used maximum intensity projection to collapse a three-dimensional z-stack into a single image. The workstation was a Dell Precision T7910 XL with an Intel^®^ Xeon^®^ Processor E5-2620 v3 (6C, 2.4GHz, 15M, 85W) and an AMD FirePro™ W7100 8GB graphics card and 32 GB random access memory (RAM). All datasets were linearly adjusted for contrast. Images were prepared for publication using Adobe Photoshop CS6 Extended, version 13.0.1 (Adobe Systems Inc, San Francisco, CA, USA).

## Supporting information

Supplementary Movies – List

## Acknowledgements

This work was made possible through a Royal Society Research Grant [RG120274], an innovation voucher from the University of Essex [DBF6000] and a Royal Society Industry Fellowship [IF150018] to PPL. PPL would like to thank Russell Smart for aquarium maintenance and Tony Jordan for production of customised parts. PPL also thanks the open-source communities OpenSPIM and μManager for support, as well as Cairn Research, 89North, Nikon Instruments UK, Alex Gardiol from Olympus Keymed UK, and Scott Young, Matt Preston and Daniel Croucher from Teledyne Photometrics for equipment loans. PPL is grateful to Loretta Roberson, Amy Gladfelter, Hari Shroff, Abhishek Kumar, Paul Maddox, Louis Kerr, Pavel Tomancak, Emmanuel Reynaud, Tali Mass, Maayan Neder, Philip M. Mullineaux, Marino Exposito-Rodriguez and Jean A. Laissue for support and critical discussions.

## Author contributions

PPL designed the study, conceived and developed the L-SPI described here. PPL, YG and CQ performed the experiments. PPL and DS interpreted the results and wrote the paper.

## REFERENCES

Brown, B. E., Letissier, M. D. A. and Dunne, R. P. (1994). Tissue retraction in the scleractinian coral Coeloseris mayeri, its effect upon coral pigmentation, and preliminary implications for heat balance. Mar. Ecol. Prog. Ser. 105, 209–218.

Caldwell, J. M., Ushijima, B., Couch, C. S. and Gates, R. D. (2017). Intra-colony disease progression induces fragmentation of coral fluorescent pigments. Sci. Rep. 7, 1–9.

Carlton, P. M., Boulanger, J., Kervrann, C., Sibarita, J.-B., Salamero, J., Gordon-Messer, S., Bressan, D., Haber, J. E., Haase, S., Shao, L., et al. (2010). Fast live simultaneous multiwavelength four-dimensional optical microscopy. Proc. Natl. Acad. Sci. U. S. A. 107, 16016–22.

Crossland, C. J. and Barnes, D. J. (1977). Gas-exchange studies with the staghorn coral Acropora acuminata and its zooxanthellae. Mar. Biol. 40, 185–194.

Edelstein, A., Amodaj, N., Hoover, K., Vale, R. and Stuurman, N. (2010). Computer control of microscopes using μManager. In Current protocols in molecular biology / edited by Frederick M. Ausubel… [et al.],.

Gladfeiter, E. H. (1982). Skeletal development in Acropora cervicornis: I. Patterns of calcium carbonate accretion in the axial corallite. Coral Reefs 1, 45–51.

Gladfelter, E. H. (1983). Skeletal development in Acropora cervicornis: II. Diel patterns of calcium carbonate accretion. Coral Reefs 2, 91–100.

Gladfelter, E. H. (2007). Skeletal development in Acropora palmata (Lamarck 1816): a scanning electron microscope (SEM) comparison demonstrating similar mechanisms of skeletal extension in axial versus encrusting growth. Coral Reefs 26, 883–892.

Gorbunov, M. Y. and Falkowski, P. G. (2002). Photoreceptors in the cnidarian hosts allow symbiotic corals to sense blue moonlight. Limnol. Oceanogr. 47, 309–315.

Gutiérrez-Heredia, L., Flood, P. M. and Reynaud, E. G. (2012). Light Sheet Fluorescence Microscopy: beyond the flatlands. In Current Microscopy Contributions to Advances in Science and Technology, pp. 838–847.

Hooton, J. W. L. (1991). Randomization tests: statistics for experimenters. Comput. Methods Programs Biomed. 35, 43–51.

Horridge, G. A. (1957). The Co-Ordination of the Protective Retraction of Coral Polyps. Philos. Trans. R. Soc. B Biol. Sci. 240, 495–528.

Huisken, J., Swoger, J., Del Bene, F., Wittbrodt, J. and Stelzer, E. (2004). Optical sectioning deep inside live embryos by selective plane illumination microscopy. Science 305, 1007–1009.

Kenkel, C. D., Traylor, M. R., Wiedenmann, J., Salih, A. and Matz, M. V (2011). Fluorescence of coral larvae predicts their settlement response to crustose coralline algae and reflects stress. Proc. R. Soc. B Biol. Sci. 278, 2691–2697.

Laissue, P. P., Alghamdi, R. A., Tomancak, P., Reynaud, E. G. and Shroff, H. (2017). Assessing phototoxicity in live fluorescence imaging. Nat. Methods 14, 657–661.

Levy, O. (2003). Photobehavior of stony corals: responses to light spectra and intensity. J. Exp. Biol. 206, 4041–4049.

Levy, O., Dubinsky, Z. and Achituv, Y. (2003). Photobehavior of stony corals: responses to light spectra and intensity. J. Exp. Biol. 206, 4041–4049.

Levy, O., Dubinsky, Z., Achituv, Y. and Erez, J. (2006). Diurnal polyp expansion behavior in stony corals may enhance carbon availability for symbionts photosynthesis. J. Exp. Mar. Bio. Ecol. 333, 1–11.

Linné, C. von (1758). Systema naturae per regna tria naturae: secundum classes, ordines, genera, species, cum characteribus, differentiis, synonymis, locis. Lugduni: Apud J. B. Delamolliere.

Mass, T., Drake, J., Haramaty, L., Rosenthal, Y., Schofield, O., Sherrell, R. and Falkowski, P. (2012). Aragonite Precipitation by “Proto-Polyps” in Coral Cell Cultures. PLoS One 7, e35049.

Mullen, A. D., Treibitz, T., Roberts, P. L. D., Kelly, E. L. A., Horwitz, R., Smith, J. E. and Jaffe, J. S. (2016). Underwater microscopy for in situ studies of benthic ecosystems. Nat. Commun. 7, 12093.

Neder, M., Laissue, P. P., Akiva, A., Akkaynak, D., Albéric, M., Spaeker, O., Politi, Y., Pinkas, I. and Mass, T. (2019). Mineral formation in the primary polyps of pocilloporoid corals. Acta Biomater. 96, 631–645.

Nielsen, D. A., Petrou, K. and Gates, R. D. (2018). Coral bleaching from a single cell perspective. ISME J. 12, 1558–1567.

Nuzzo, R. L. (2017). Randomization Test: An Alternative Analysis for the Difference of Two Means. PM&R 9, 306–310.

Ohno, Y., Iguchi, A., Shinzato, C., Gushi, M., Inoue, M., Suzuki, A., Sakai, K. and Nakamura, T. (2017). Calcification process dynamics in coral primary polyps as observed using a calcein incubation method. Biochem. Biophys. Reports 9, 289–294.

Pitrone, P. G., Schindelin, J., Stuyvenberg, L., Preibisch, S., Weber, M., Eliceiri, K. W., Huisken, J. and Tomancak, P. (2013). OpenSPIM: an open-access light-sheet microscopy platform. Nat. Methods 10, 598–9.

Renema, W., Pandolfi, J. M., Kiessling, W., Bosellini, F. R., Klaus, J. S., Korpanty, C., Rosen, B. R., Santodomingo, N., Wallace, C. C., Webster, J. M., et al. (2016). Are coral reefs victims of their own past success? Sci. Adv. 2, e1500850.

Reynaud-Vaganay, S., Gattuso, J. P., Cuif, J. P., Jaubert, J. and Juillet-Leclerc, A. (1999). A novel culture technique for scleractinian corals: application to investigate changes in skeletal d18O as a function of temperature. Mar. Ecol. Prog. Ser. 180, 121–130.

Reynaud, E. G., Krzic, U., Greger, K. and Stelzer, E. H. K. (2008). Light sheet-based fluorescence microscopy: more dimensions, more photons, and less photodamage. HFSP J. 2, 266–275.

Salih, A., Larkum, A., Cox, G., Kuhl, M. and Hoegh-Guldberg, O. (2000). Fluorescent pigments in corals are photoprotective. Nature 408, 850–853.

Schindelin, J., Arganda-Carreras, I., Frise, E., Kaynig, V., Longair, M., Pietzsch, T., Preibisch, S., Rueden, C., Saalfeld, S., Schmid, B., et al. (2012). Fiji: an open source platform for biological image analysis. Nat. Methods 9, 676–682.

Sebens, K. P. and DeRiemer, K. (1977). Diel cycles of expansion and contraction in coral reef anthozoans. Mar. Biol. 43, 247–256.

Shafir, S., Rijn, J. V. A. N. and Rinkevich, B. (2001). Nubbing of coral colonies: A novel approach for the development of inland broodstocks. Aquarium Sci. Conserv. 3, 183–190.

Shafir, S., Van Rijn, J. and Rinkevich, B. (2003). The use of coral nubbins in coral reef ecotoxicology testing. Biomol. Eng. 20, 401–406.

Shapiro, O. H., Kramarsky-Winter, E., Gavish, A. R., Stocker, R. and Vardi, A. (2016). A coral-on-a-chip microfluidic platform enabling live-imaging microscopy of reef-building corals. Nat. Commun. 7, 10860.

Stuurman, N. and Swedlow, J. R. (2012). Software tools, data structures, and interfaces for microscope imaging. Cold Spring Harb. Protoc. 2012, 50–61.

Swain, T. D., Schellinger, J. L., Strimaitis, A. M. and Reuter, K. E. (2015). Evolution of anthozoan polyp retraction mechanisms: Convergent functional morphology and evolutionary allometry of the marginal musculature in order Zoanthidea (Cnidaria: Anthozoa: Hexacorallia). BMC Evol. Biol. 15, 1–19.

Sweatman, H., Delean, S. and Syms, C. (2011). Assessing loss of coral cover on Australia’s Great Barrier Reef over two decades, with implications for longer-term trends. Coral Reefs 30, 521–531.

Venn, A. A., Tambutté, E., Lotto, S., Zoccola, D., Allemand, D. and Tambutté, S. (2009). Imaging intracellular pH in a reef coral and symbiotic anemone. Proc. Natl. Acad. Sci. U. S. A. 106, 16574–16579.

Venn, A., Tambutté, E., Holcomb, M., Allemand, D. and Tambutté, S. (2011). Live tissue imaging shows reef corals elevate pH under their calcifying tissue relative to seawater. PLoS One 6, e20013.

Venn, A. A., Tambutté, E., Holcomb, M., Laurent, J., Allemand, D. and Tambutté, S. (2013). Impact of seawater acidification on pH at the tissue-skeleton interface and calcification in reef corals. Proc. Natl. Acad. Sci. U. S. A. 110, 1634–9.

