## Supplementary Movies – List for "Light-induced polyp retraction and tissue rupture in the photosensitive, reef-building coral *Acropora muricata*"

**Supplementary Movies – List with descriptions**

| Title | Corresponding figure | Description |
|---|---|---|
| Supplementary Movie S1 | Fig. 2 C | Time-lapse movie of small polyp over 4.7 h |
| Supplementary Movie S2 | Fig. 3 A, top row | Polyp emergence in low irradiance conditions (18.4 $mW/cm^2$) over 3h |
| Supplementary Movie S3 | Fig. 3 A, bottom row | Polyp emergence in high irradiance conditions (82.6 $mW/cm^2$) |
| Supplementary Movie S4 | Fig. 3 B | Polyp switching from low to high irradiation. The polyp emerges at low irradiance (9.2 $mW/cm^2$) and retracts at high irradiance (59.7 $mW/cm^2$) |
| Supplementary Movie S5 | Fig. 4 A, B | Polyp, coral tissue and zooxanthellae in low light (9.2 $mW/cm^2$) over 6h |
| Supplementary Movie S6 | Fig. 4 C | Close-up of zooxanthellae from supplementary movie S5 |
| Supplementary Movie S7 | Fig. 5 C | Zooxanthellae in low light (9.2 $mW/cm^2$) over 7h, then 90 min in high light (59.7 $mW/cm^2$)<br>*Note: The movie corresponding to Fig. 5 A, B and showing polyp, coral tissue and zooxanthellae in the full field of view is available at: https://www.nikonsmallworld.com/galleries/small-world-in-motion* |
| Supplementary Movie S8 | Fig. 6 | Tissue rupture in high light (59.7 $mW/cm^2$) after 9h |
| Supplementary Movie S9 | Fig. 7 A | Retracted polyps at the tip of a coral branch, imaged with CLSM |
| Supplementary Movie S10 | Fig. 7 C | Polyp imaged over 6h with WFM |
