## Supplementary Table 1 for "Light-induced polyp retraction and tissue rupture in the photosensitive, reef-building coral *Acropora muricata*"

***Supplementary Table 1*** *Lateral resolutions and light-sheet dimensions*

*a) Lateral resolutions using different detection objectives*

| ***Magnification, type*** | *Numerical aperture* | *Theoretical Rayleigh resolution limit (µm, at 510 nm)* | *Measured in thick samples (µm)* | *Manu-facturer* | *Objective model* |
|---|---|---|---|---|---|
| 4x, air | 0.1 | 3.1 | 4.1 ± 0.8 | Olympus | PLAN 4X |
| 10x, water immersion | 0.3 | 1.0 | 1.4 ± 0.2 | Olympus | UMPLFLN 10XW |
| 20x, water immersion | 0.5 | 0.6 | 0.7 ± 0.1 | Olympus | UMPLFLN 20XW |
| 40x, water immersion | 0.8 | 0.4 | 0.6 ± 0.1 | Olympus | LUMPLFLN 40XW |

*b) Light-sheet dimensions using different cylindrical lenses*

| *Focal length (mm)* | *Measured waist (µm, FWHM)* | *confocal parameter (mm)* | *Maximum width (mm)* | *Manufacturer* | *Application* |
|---|---|---|---|---|---|
| 50 | 20.7 ± 0.8 | 1.6 | 20 | Thorlabs | For large FOVs |
| 30 | 11.7 ± 0.8 | 0.6 | 20 | Thorlabs | For single polyps |
