## Supplementary Table 2 for "Light-induced polyp retraction and tissue rupture in the photosensitive, reef-building coral *Acropora muricata*"

*Powers, irradiances and experimental durations used in this study.*

| Technique | Power [µW] | Detection objective magni-fication | Irradiance [$mW/cm^2$] | Photon flux [µmol $m^{-2}$ $s^{-1}$] | Dura-tion [h] | Mode | Physiological effect |
|---|---|---|---|---|---|---|---|
| L-SPI (non-invasive) | 10 | 0.5x – 40x | 9.2 | 375 | 8.3 ± 2.0 | Continuous image acquisition | Polyps emerge; no long-term effects measured; slight photo-bleaching of coral tissue |
| L-SPI (invasive) | 52 - 90 | 10x | 48-85 | 1948-3372 | 7 - 9 | Continuous image acquisition | Polyps retract; pronounced photo-bleaching; surge of auto-fluorescence; rupture of coral tissue |
| WFM | 170 | 4x | 6.2 | 253 | 6 | 5 min intervals | Polyps emerge; pronounced photo-bleaching of coral tissue |
| CLSM | 1.1 | 10x | 34’414 | $1.4 * 10^6$ | 3 | 1 h intervals | Retraction of polyps and coral tissue; long-term damage leads to tissue rupture |
