## Supplementary Table 3 for "Light-induced polyp retraction and tissue rupture in the photosensitive, reef-building coral *Acropora muricata*"

| Components list | | | | |
|---|---|---|---|---|
| ***Optomechanical components:*** | **Product code** | **Quantity** | **Provider** | **Comment** |
| Postholders | UPH1.5 | 8 | Thorlabs | |
| Posts | TRP1.5-P5 | 2 | Thorlabs | |
| Kinematic mirror mount | POLARIS-K1 | 2 | Thorlabs | placed before cylindrical lenses |
| 1" Full Gimbal Mount | GMB1 | 1 | Thorlabs | placed after beam expander and before rotating mirror for alignment control |
| Broadband Dielectric Mirror | BB1-E02 | 3 | Thorlabs | |
| Lens mount for 1/2" optics | LMR05 | 2 | Thorlabs | beam expander |
| Achromatic doublet f=25mm | AC127-025-A-ML | 1 | Thorlabs | beam expander |
| Achromatic doublet f=50mm | AC127-050-A-ML | 1 | Thorlabs | beam expander |
| Adjustable Mechanical Slit, Metric | VA100/M | 1 | Thorlabs | place after rotating mirror for varying width of light sheet |
| Kinematic Rectangular Optic Mount, Right Handed, Adjustable Height | KM100C | 1 | Thorlabs | |
| Kinematic Rectangular Optic Mount, Left Handed, Adjustable Height | KM100CL | 1 | Thorlabs | |
| Small Adjustable Clamping Arm, 6-32 Threaded Post | PM3 | 2 | Thorlabs | |
| N-BK7 Plano-Convex Cylindrical Lens, f = 50.00 mm, H = 20.00 mm, L = 22.0 mm, Antireflection Coating: 350-700 nm | LJ1821L1-A | 2 | Thorlabs | For light-sheet dimensions, see Supplementary Table 1 |
| N-BK7 Plano-Convex Cylindrical Lens, f = 30.00 mm, H = 20.00 mm, L = 22.0 mm, Antireflection Coating: 350-700 nm | LJ1212L1-A | 2 | Thorlabs | The shorter focal length of these cylindrical lenses requires that the kinematic mounts (KM100C and KM100L) are moved closer to the sample |
| Pedestal Pillar Post | RS1.5P | 6 | Thorlabs | 4 for mounting heat sink and laser, 2 for z-stage |
| Small clamping fork | CF125 | 6 | Thorlabs | 4 for mounting heat sink and laser, 2 for z-stage |
| Ø1.5" Mounting Post Bracket | C1505 | 1 | Thorlabs | for mounting microscope pillar (base removed) |
| 50:50 Non-Polarizing Beamsplitter Cube, 400 - 700 nm, 5 mm | BS007 | 1 | Thorlabs | |
| Right-Angle Prism Dielectric Mirror, 400 - 750 nm, L = 10.0 mm | MRA10-E02 | 1 | Thorlabs | |
| Ø1", SM1-Mounted N-BK7 Ground Glass Diffuser, 600 Grit | DG10-600-MD | 1 | Thorlabs | for brightfield imaging |
| Right-Angle Prism Dielectric Mirror, 400 - 750 nm, L = 25.0 mm | MRA25-E02 | 1 | Thorlabs | for brightfield imaging; position underneath hole of L-shaped bracket fitted to z-stage |
| ***Z-stage:*** | | | | |
| Single-Axis, 0.98" Travel, Motorized Translation Stage | PT1-Z8 | 1 | Thorlabs | |
| PT-Series Angle Bracket | PT102/M | 1 | Thorlabs | |
| T-Cube DC Servo Motor Controller (Power Supply Not Included) | TDC001 | 1 | Thorlabs | |
| 15 V Power Supply Unit for a Single T-Cube | TPS001 | 1 | Thorlabs | |
| Motor Extension Cable, 2.5 m, DB15 Male to DB15 Female | PAA632 | 1 | Thorlabs | |
| ***Laser:*** | | | | |
| OBIS 488 nm LS 100 mW | OBIS 488 LS | 1 | Coherent, Inc. | |
| ***Rotating mirror:*** | | | | |
| Panasonic AN8248NSB | n/a | 1 | Panasonic Corp. | See weblink below |
| 375W Linear DC Variable Voltage Bench Power Supply | RP10L | 1 | Maplin | |
| ***Accessories:*** | | | | |
| 8-32 Cap Screw and Hardware Kit | HW-KIT1 | 1 | Thorlabs | |
| 1/4"-20 Cap Screw and Hardware Kit | HW-KIT2 | 1 | Thorlabs | |

| | | | | |
|---|---|---|---|---|
| Laser Safety Glasses | LG3 | 1 | Thorlabs | |
| USB Temperature and Humidity Data Logger, -50 °C to 150 °C | TSP01 | 1 | Thorlabs | |
| Additional External Temperature Probe, -15 °C to 200 °C | TSP-TH | 1 | Thorlabs | |
| Plain Glass Slides 76 mm x 39 mm x 1.0-1.2 mm (Pack of 100) | AGL4222A | 1 | Agar Scientific, UK | for observation vessel |
| Large Glass Slides 102 mm x 83 mm (Pack of 36) | AGL4380-2 | 1 | Agar Scientific, UK | for observation vessel |
| AquaMate Silicone Sealant, fungicide and solvent-free | AQUATR | 1 | Everbuild, UK | for observation vessel |
| Monument Tools Fluorescein Drain Dye | 31595 | 1 | Screwfix | for lightsheet measurements |

Rotating mirror from laser printer scanner assembly, available at:
https://www.itcsales.co.uk/cgi-bin/sh000002.pl?WD=scanner&PN=HP-LaserJet-1300-1150-3380-Scanner-Assembly-RM1-0524-7650%2ehtml#SID=473

Online instructions for controlling OBIS laser using Arduino and μManager:
https://actin.cn/2014/12/Arduino-UNO-control-OBIS-via-digital-trigger
https://micro-manager.org/wiki/Control_laser_shutters_with_Arduino
